# Task-Dependent Modulation of Upper-Extremity Muscle Synergies during Object Slippage

**DOI:** 10.64898/2026.09.10.746922

**Authors:** Ayesha Tooba Khan, Biswarup Mukherjee, Deepak Joshi

## Abstract

Preventing object slippage requires the central nervous system to control and coordinate the muscle groups while adapting to the variations in the slip characteristics. Our study investigates how slip characteristics, such as slip direction, slip distance, and slip speed, affect the modulation of muscle synergies as a manifestation of object slippage. Our results reveal that there are shared synergies existing in the upward and downward slip emulation, and that these shared synergies are dependent on slip characteristics. However, there were no specific synergies found to exist in the upward and downward slip conditions. Our results also indicate the existence of combined synergies, as the motor modules in the upward slip direction were found to have shared synergies with multiple motor modules in the downward slip direction. We also observed that the number of shared synergies varied depending on the slip distance and slip speed. Together, these findings suggest that the CNS flexibly reconfigures a limited set of muscle synergies to accommodate varying slip dynamics, balancing control stability with adaptability.

## I. Introduction

The intricate hand movements required to perform activities of daily living are a remarkable outcome of the robust bidirectional coordination between the central nervous system (CNS) and the peripheral nervous system [1]. With approximately 68 muscles in the upper limb, the challenge of understanding the neuroscience of hand dexterity revolves around a fundamental question: how does the CNS manage the enormous complexity of a musculoskeletal system having multiple degrees of freedom? One of the leading hypotheses to decode the complex neuromuscular coordination is the muscle synergy framework, where spatial and temporal coherent activations of a group of muscles are believed to be innervated by a single neural command, referred to as synergies [2]. Muscle synergies enable translation of higher-level motor intentions into low-level muscle activation patterns [3]. Human movement-related studies have shown that typically 4-5 synergies are required to explain most of the muscle activation patterns in healthy individuals [4]. However, this dimensional collapse does not stem from the biomechanics of the musculoskeletal system itself but rather emerges from the neural control strategies governing the movement [5]. Given its ability to simplify the complexity of neuromuscular control, the synergy framework has found broad practical relevance from clinical rehabilitation to skilled motor performance [6]– [8].

In recent years, muscle synergies have been thoroughly investigated to understand the functional connectivity of the synergistic patterns in individuals with stroke [9], cerebral palsy [10], cerebellar ataxia [11], and other neuromuscular disorders. The concept of muscle synergies provides a promising framework for characterizing neuromuscular organization for various sensorimotor deficits and, therefore, possesses great potential in developing rehabilitation strategies [12]– [14]. A recent study reported the development of a new intermuscular coordination pattern following alterations to the existing muscle synergies through short-term training [15]. Furthermore, several sports research studies have also utilized the concept of muscle synergies to assess the performance of athletes and to develop the strategic planning of training programs for players [16]–[18]

While these studies demonstrate the wide applicability of the muscle synergy framework, understanding the mechanistic modulation of synergies under varying movement conditions remains equally crucial. The experimental evidence suggests that reflex and voluntary mechanisms often recruit the same motor modules; however, the motor primitives often exhibit variations during movement [19]. The speed of hand movement [20], weight support [21], and weight lifting [22] were reported to have an influence on motor primitives, while the effect of body position under the influence of gravity was found to affect the motor modules [23]. There is an extensive literature available to support altered motor movement under the effect of gravity, which suggests the ability of the CNS in developing the optimal motor control strategies [24], [25]. One of the most practical scenarios to observe the coordination of hand movement under the gravitational effect is the event of object slippage. Mechanical stimuli, presented either as perturbations to the shoulder muscles or slip sensations at the fingertips, require the rapid integration of somatosensory feedback and may impose different neuromechanical demands depending upon the external stimuli [26]–[29], which may lead to changes in muscle synergies. However, despite extensive evidence on gravity-dependent modulation of synergies, limited attention has been given to how slip-related mechanical stimuli influence these coordination patterns. Therefore, it is crucial to investigate how the control architecture of the CNS adapts to different slip characteristics. Specifically, it remains unclear whether this adaptation involves a reorganization of underlying muscle coordination patterns or a modulation of the recruitment of existing modules.

Our research thus investigates the interactive effect of slip characteristics on muscle synergies during slip emulation. We hypothesized that there would be shared synergies between upward and downward slip conditions; however, we were keen to see whether these synergies are altered by variations in slip distance and slip speed. Variations in slip distance and slip speed may force the CNS to recruit shared synergies differently or to develop new or reorganized synergies to counterbalance the mechanical disturbance. Therefore, we also investigated whether the number of shared synergies varies with the slip distance and slip speed. To our knowledge, this is the first study to investigate muscle synergies for such a fine-movement task, with an emphasis on in-depth analysis of their modulation across varying slip characteristics, including direction, distance, and speed.

## II. Materials and Methods

### A. Experimental Setup and Paradigm

A detailed overview of the device and experiment protocol (Refer Fig. 1(a)) has been published in our previous research works, where psychophysics [30] and force dynamics [26] of the object slippage based on slip characteristics were investigated. Briefly, a Slip Inducing Device (SID) with a control unit driven by a microcontroller (Teensy 3.2, PJRC Inc., USA) was developed to simulate the artificial sense of object slippage. The device, equipped with a force measurement unit comprising the four load cells (3 Kg, YZC-131, China) to measure the grip and tangential forces, was capable of simulating the slip in both upward and downward directions with multiple speeds and distances (Refer Fig. 1(b)). A MATLAB app-based user interface (MATLAB 2023b, The MathWorks Inc., USA) was developed to facilitate the data acquisition process by establishing the communication between the control unit of the SID, the EMG data acquisition system (PLUX Biosignals, Lisbon, Portugal), and the PC (Windows 10, 64-bit, Core i7 CPU, 3.2 GHz processor with 16 GB RAM).

**Fig. 1.**
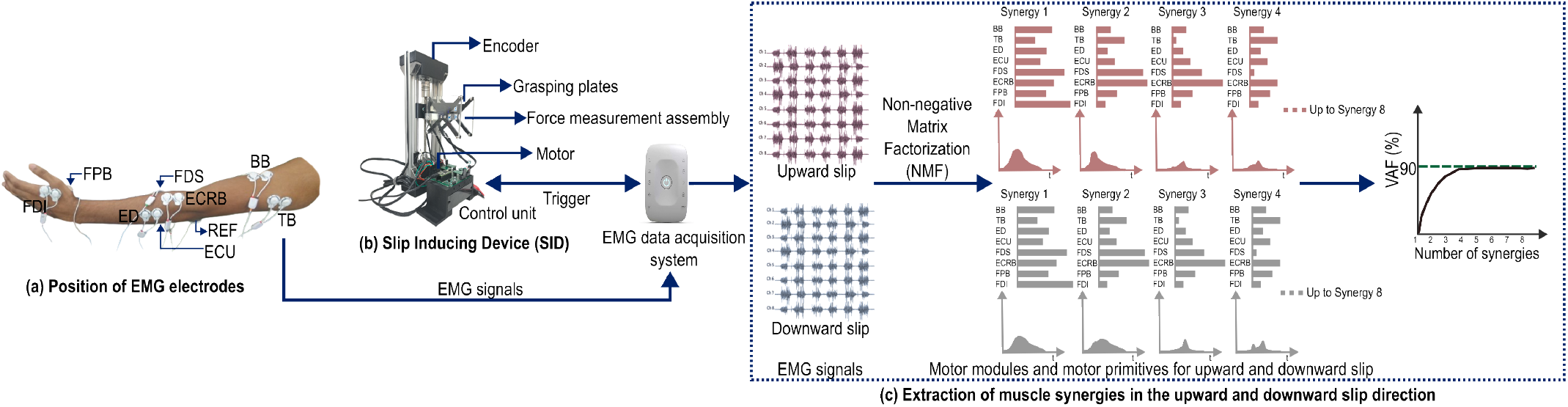
(a) Position of the EMG electrodes on the upper extremity. (b) The Slip Induction Device SID shows the force measurement assembly, grasping plates, and the control unit. (c) Extraction of muscle synergies from the EMG data during up and down slip directions.

The participant was instructed to hold onto the grasping plates using a prehensile grip with the dominant hand while seated in an adjustable chair. During the experiment, the dominant hand was positioned on the armrest such that the angle between the long axes of the humerus and the radius in the sagittal plane was maintained at 135^*°*^. The auditory and the visual cues arising due to the device operation were occluded by playing the white noise in the active noise cancellation headphones (Bose QuietComfort 25, Bose Corp., USA) and by putting a black curtain alongside the device, respectively.

Each participant performed four blocks of the experiment. Each block had a distinct command sequence for slip stimuli comprising a combination of a particular slip distance and slip speed alternately in the upward and downward slip directions. We considered five slip distances (2 mm to 10 mm with a linear increment of 2 mm) and slip speeds (2 mm/s to 10 mm/s with a linear increment of 2 mm/s), which were validated to produce slip sensations in our previous psychophysical study [30]. Thus, each block had fifty unique stimuli (2 slip direction × 5 slip distance × 5 slip speed). We also interleaved six catch trials in the command sequence to minimize the motor learning effects. Thus, in total, there were 56 trials in each block. At the beginning of each block, a ‘start trigger’ was sent by the SID control unit to the trigger module of the EMG data acquisition to stream a time-synchronized EMG activity. During each trial, the participant was instructed to hold onto the grasping plates for at least 125 ms, maintaining the grip force with the fingers within 1.5±0.1 N. This was regarded as the calibration phase of the trial, during which the participant was provided with the visual feedback of the instantaneous grip force. The visual feedback of the instantaneous grip force exerted by the fingers on the grasping plates would disappear once the calibration is achieved. The participant had a maximum of 6 s to complete the calibration, failing which the slip stimulus for that particular trial would not be presented to the participant, and it would be deemed a failed trial. Alternatively, if the calibration succeeds, the trial would then proceed to the slip simulation phase, where an artificial slip would be simulated in a particular slip direction with a specified slip distance and slip speed as per the command sent by the control unit of SID. The SID control unit was programmed to send the digital markers to segregate the calibration and the slip simulation phases. After each block, the participant was provided a rest of 10 min to prevent muscle fatigue. The total duration of the study was approximately 180 min.

### B. Participants

Eight non-disabled volunteers (two females and six males; aged 27.25±4.83) provided written informed consent to participate in the study. The recruited participants had no history of contact dermatitis, neuromuscular disorder, or surgical intervention in the upper extremity. All participants reported having either normal or corrected-to-normal vision and being right-hand dominant. The experimental protocols were reviewed and approved by the Institute Ethics Committee at the Indian Institute of Technology Delhi (Ref no: 2021/P052).

### C. Data Collection

The surface electromyography (EMG) data were recorded using an eight-channel EMG system (PLUX Biosignals, Lisbon, Portugal). The physiological signals were transmitted to the PC via Bluetooth using a wireless eight-channel biosignalsplux kit with a sampling rate of 1000 Hz and 16-bit resolution per channel. The OpenSignals v2.2.5 software (PLUX Biosignals, Lisbon, Portugal) was used for real-time data acquisition. The auxiliary port of the wireless hub was used to synchronize the EMG data with the operation of the Slip Inducing Device (SID) and the MATLAB GUI, utilizing a digital trigger signal generated by the control unit of the SID. First, the skin was cleaned and abraded using the skin preparation gel (Nuprep, PLUX Biosignals, Lisbon, Portugal). Then, eight gelled self-adhesive disposable Ag/AgCl electrodes (PLUX Biosignals, Lisbon, Portugal) with bipolar configuration were placed on the muscle belly of the upper arm muscles (biceps brachii (BB) (long head), triceps brachii (TB) (lateral head)), forearm muscles (extensor digitorum (ED), extensor carpi ulnaris (ECU), flexor digitorum superficialis (FDS), extensor carpi radialis brevis (ECRB)), and hand muscles (first dorsal interosseous (FDI), flexor pollicis brevis (FPB)) as shown in Fig. 1(a). The reference electrode (REF) was placed at the Olecranon.

### D. Data Analysis

#### 1) Data Preprocessing

The acquired continuous EMG signals were segmented for each trial using the markers for the calibration and the slip simulation phase. A 2nd-order Butterworth zero-phase digital filter with cutoff frequencies of 10-450 Hz was used to bandpass filter the segmented EMG data. The upper root mean square envelope was extracted from the filtered EMG signals using a sliding window of 125 ms. The extracted EMG envelopes from the slip simulation phase were corrected for baseline by subtracting the average value of the 125 ms of data preceding the slip onset in the calibration phase. The baseline-corrected envelopes for each channel were normalized for each subject, considering the maximum and minimum values across all trials and blocks [31].

#### 2) Muscle Synergies Extraction

The non-negative matrix factorization (NMF) was used to extract muscle synergies from the preprocessed EMG envelopes. We considered only the regular trials (excluding six catch trials) for synergy extraction corresponding to 25 upward slip and 25 downward slip conditions for each subject (Refer Fig. 1(c)). The preprocessed EMG envelopes for each stimulus were decomposed into a set of muscle synergies (motor modules) and their activation coefficients (motor primitives) as:

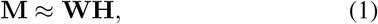

where **M** ∈ ℝ^*m×t*^ represents the EMG data matrix with *m* muscles and *t* time samples, **W** ∈ ℝ^*m×r*^ is the muscle weight matrix containing *r* synergies, and **H** ∈ ℝ^*r×t*^ is the activation coefficient matrix describing the temporal contribution of each synergy.

The decomposition is achieved by minimizing the reconstruction error based on the Frobenius norm:

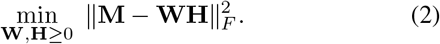

After the extraction of the muscle synergies, we considered Variance Accounted For (VAF) as the metric to determine the minimum number of synergies that could account for at least 90 % of the variability of the EMG. Mathematically, VAF is computed as:

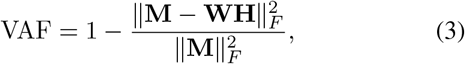

where ∥ · ∥_*F*_ denotes the Frobenius norm. A higher VAF indicates that the extracted synergies effectively capture the original EMG variability.

Thus, the choice of the number of synergies considered to investigate the shared synergies between upward and downward slip conditions was governed by the minimum number of synergies that could account for at least 90 % of the variability of the EMG. After NMF and computation of VAF, we found that at least four synergies would be required to explain 90 % variance in the upward and downward slip emulation. Therefore, the motor modules from four synergies were considered for further investigation of muscle synergies shared between upward and downward slip directions.

#### 3) Reordering of Muscle Synergies

The NMF does not retain the order of synergies across blocks and subjects; therefore, it is crucial to sort the extracted synergies based on their similarity to one another across the blocks and subjects. Thus, after extracting synergies for each block from all subjects, the synergies were reordered to group similar synergies across blocks and subjects. For this, the motor modules, comprising eight muscles corresponding to four synergies, were concatenated across all four blocks for each of the eight subjects, resulting in an array of 8×4×32 as shown in Fig. 2. Then, these arrays were pooled for 25 upward and downward slip conditions, and the k-means clustering algorithm was used to generate the canonical motor modules (8×4) for upward and downward slip conditions, separately. The cosine similarity (4×4) between the unordered motor modules and the canonical motor module for upward and downward slip conditions was computed as:

**Fig. 2.**
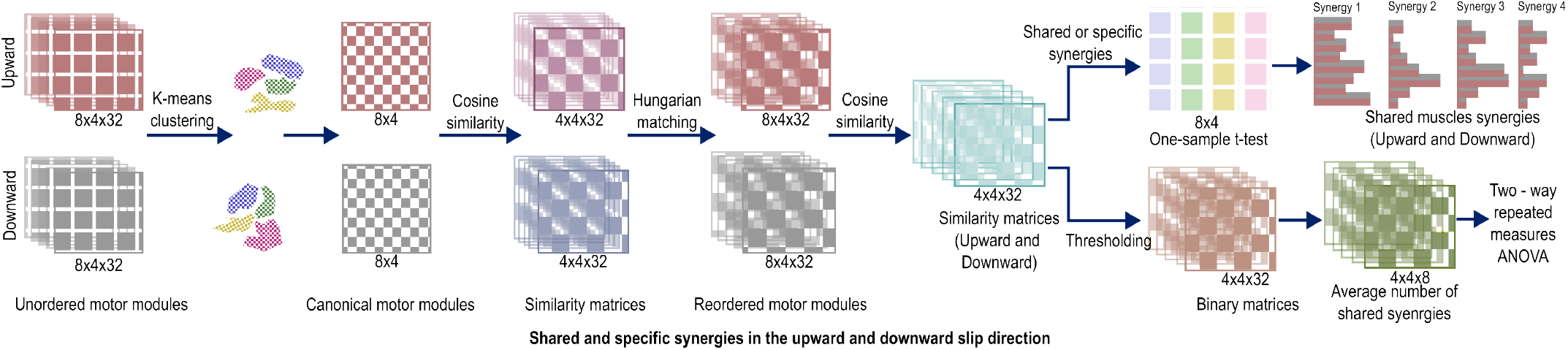
Schematic diagram showing the process of reordering of the muscle synergies and the determination of shared and specific synergies for a particular slip distance and speed in upward and downward slip directions.

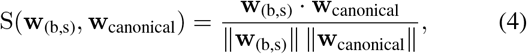

where **w**_(b,s)_ represents the motor module of subject *s* during block *b*. Finally, Hungarian matching was used to reorder the similar synergies across all the participants for both slip directions.

#### 4) Shared Muscle Synergies

These reordered motor modules were then used to compute the cosine similarity between up and down slip directions for multiple slip distances and speeds (Fig. 2). The similarity between corresponding muscle synergies extracted during upward and downward slip conditions for each slip distance and speed was quantified using the cosine similarity as:

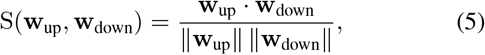

where S(**w**_up_, **w**_down_) ranges from 0 (orthogonal) to 1 (parallel), indicating the degree of shared structure between the synergies under the two slip directions. This resulted in an array of 4×4×32 for 25 slip conditions (5 distances ×5 speeds).

The cosine similarity values were averaged across blocks for each subject to determine statistically significant thresholds (discussed in the next subsection) for categorizing muscle synergies as shared and specific. A cosine similarity threshold of 0.8 was considered to categorize the muscle synergies as shared, while a threshold of 0.4 was used to designate the muscle synergies as specific. Furthermore, we performed thresholding using the statistically significant threshold to compute the number of shared synergies (*N*_*shared*_) between upward and downward slip conditions for each slip distance and speed. The *N*_*shared*_ corresponding to different slip distances and speeds was averaged across the blocks for each subject. This was done to investigate the effect of varying slip distance and slip speeds on the average number of shared synergies.

#### 5) Statistical Analysis

A one-sample t-test on average cosine similarity values for each slip distance and speed was performed for all four synergies to determine whether the motor module was significantly shared (right-tailed one-sample t-test) or was significantly specific (left-tailed one-sample t-test)between the upward and downward slip directions. The threshold values of 0.8 and 0.4 were used to test for shared and specific synergies, respectively. Additionally, a two-way repeated measures ANOVA was performed to determine if there was a significant interaction between slip distance and slip speed affecting the average number of shared synergies. All statistical analyses were performed with a significance level of *α* = 0.05. We also determined the effect size and statistical power of the study for every statistically significant outcome. The statistical analysis was performed in IBM SPSS Statistics (Version 28.0, IBM Corp, Armonk, USA).

## III. Results

The NMF was performed on normalized EMG envelopes to extract the muscle synergies. The VAF of 90 % was obtained corresponding to four synrgies. The motor modules, as well as motor primitives, for all four synergies were reordered and averaged across all the participants. Fig. 3 shows the EMG envelopes for the upward and downward slip conditions and the associated motor modules and motor primitives corresponding to the four synergies for a slip distance of 10 mm and slip speed of 2 mm/s. The motor modules across four synergies show differences in muscle weights for the upward and downward directions. Moreover, the motor primitives across four synergies also exhibit direction-based variation, which is more pronounced corresponding to the early voluntary phase (0.3 s approximately) in synergy 2 and at the slip onset time for synergy 3.

**Fig. 3.**
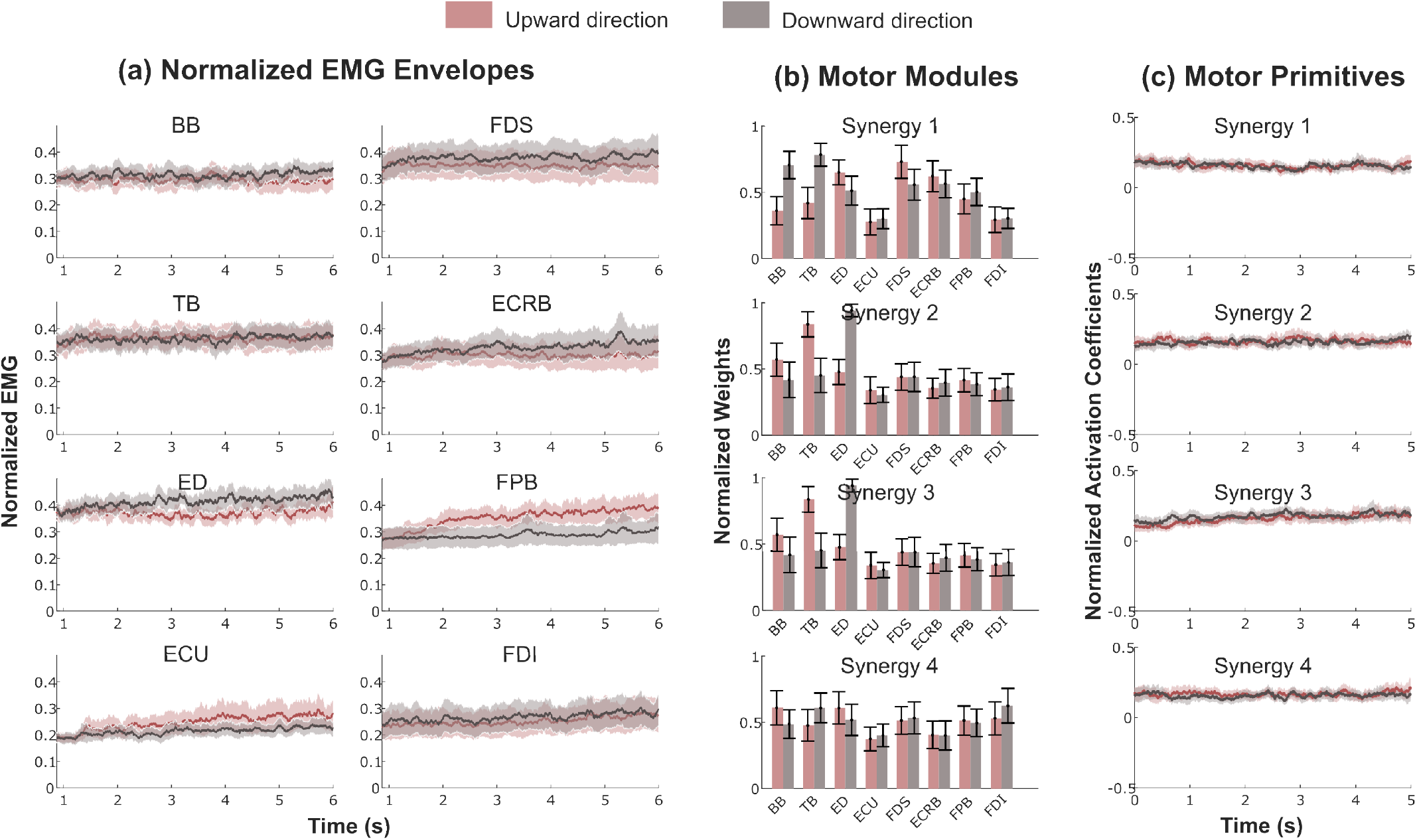
(a) Normalized EMG envelopes from the eight-channel recorded EMG. (b) The normalized weights of the motor modules corresponding to eight muscles for four synergies. (c) The normalized activation coefficients of the motor primitives for four synergies.

The average motor modules and the motor primitives corresponding to four synergies for all combinations of slip distance and slip speed in the upward and downward slip direction are shown in the supplementary material (see Fig. S1 to Fig. S8).

### A. Existence of shared and specific synergies in Upward and Downward Slippage

The right-tailed one-sample t-test revealed that synergies were significantly shared between the up and down slip conditions for specific combinations of slip distance and speed(Refer Fig. 4). Synergies were primarily shared at smaller to medium slip distances and predominantly at medium to higher slip speeds. Notably, most of the significantly shared synergies across slip conditions originated from the motor modules corresponding to synergies 1 and 4, whereas synergies 2 and 3 did not exhibit consistent sharing across subjects. In contrast, the left-tailed one-sample t-test did not reveal statistically significant evidence for the presence of specific synergies between upward and downward slip conditions.

**Fig. 4.**
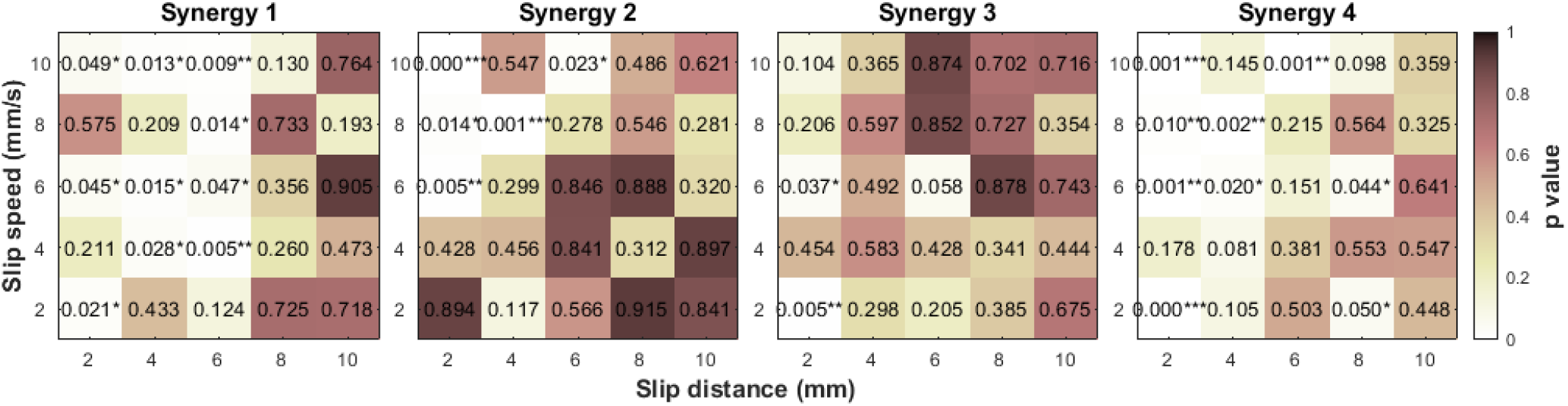
(a) The results of the one-sample t-test to determine the shared synergies for each synergy module. The significantly shared synergies across different slip distances and speeds are marked with an asterisk.

### B. Modulation of number of shared synergies based on slip distances and slip speeds

As our results suggested that the existence of shared synergies was driven by the choice of slip distances and slip speeds, we were keen to further see if *N*_*shared*_ is also modulated by the characteristics of the mechanical slip stimuli. We determined the effect of varying slip distances and slip speeds on the number of uniquely shared synergies, i.e., synergies from upward slip that are shared with the corresponding synergies from downward slip. Although we did not find any significant two-way interaction between slip distance and slip speed modulating *N*_*shared*_, we found that the slip distance alone exhibited a significant effect on *N*_*shared*_ between up and down slip conditions for all uniquely shared synergies (Refer Table I). In addition to the slip distance, the number of shared synergies for synergy 2 also varied based on slip speed (*F* (4, 28) = 3.972, *p* = 0.011).

**TABLE I.**
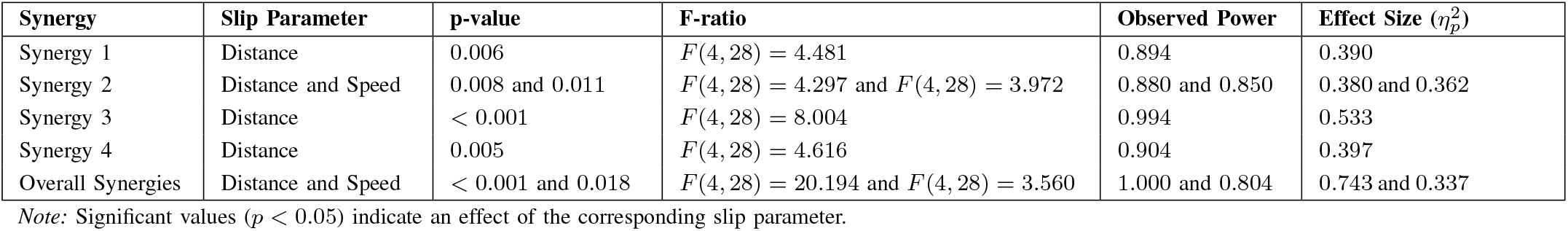
Effect of interaction of slip parameters on the number of shared synergies (*N*_*shared*_)

| Synergy | Slip Parameter | p-value | F-ratio | Observed Power | Effect Size ($\eta_p^2$) |
| --- | --- | --- | --- | --- | --- |
| Synergy 1 | Distance | 0.006 | $F(4, 28) = 4.481$ | 0.894 | 0.390 |
| Synergy 2 | Distance and Speed | 0.008 and 0.011 | $F(4, 28) = 4.297$ and $F(4, 28) = 3.972$ | 0.880 and 0.850 | 0.380 and 0.362 |
| Synergy 3 | Distance | $< 0.001$ | $F(4, 28) = 8.004$ | 0.994 | 0.533 |
| Synergy 4 | Distance | 0.005 | $F(4, 28) = 4.616$ | 0.904 | 0.397 |
| Overall Synergies | Distance and Speed | $< 0.001$ and 0.018 | $F(4, 28) = 20.194$ and $F(4, 28) = 3.560$ | 1.000 and 0.804 | 0.743 and 0.337 |

Note: Significant values ( $p < 0.05$ ) indicate an effect of the corresponding slip parameter.

We also observed that, in addition to those uniquely shared synergies between up and down slip conditions, there are several other pairs of shared synergies as well. For instance, synergy 1 of upward slip was found to be uniquely shared with synergy 1 of downward slip, as well as synergy 3 of downward slip. Therefore, we also tested whether there exist any two-way interactions of slip distance and slip speed on the overall number of shared synergies. We found that the slip distance and slip speed modulate the overall number of shared synergies; however, no significant two-way interaction was found between slip distance and slip speed, as mentioned in Table I.

## IV. Discussion

In this study, we investigated the effect of slip characteristics, i.e., direction, distance, and speed, on muscle synergies resulting from the emulation of the vertical slip. The 8-channel EMG signals were recorded from healthy participants while they experienced the mechanical stimuli at the fingertips during prehensile grip. The main objectives of the study were to investigate the existence of shared synergies between upward and downward slip conditions and their modulation based on slip distance and slip speed.

### A. Slip direction exhibits modulation of muscle synergies

Our results revealed direction-based differences in the motor primitives as well as the motor modules; however, these variations were not prominent, except in some cases for FDS and ED. Interestingly, these variations also seemed to be dependent on slip distance and slip speed. The observed variations in the motor modules, especially for synergy 1 and synergy 2, suggest adaptive changes in the activated group of muscles, while the alterations in the motor primitives represent the temporal dynamics of the upward and downward slips. It is more likely to have direction-based variations in motor primitives rather than in the motor modules, as it is the stimulus dynamics that alter the neural mechanisms while the participant tries to maintain a stable grasp upon experiencing the slip. This may have led to the existence of the direction-based shared synergies.

### B. Shared synergies and their modulation based on slip distance and speed

The existence of shared synergies between the motor movements for upward and downward slips suggests that functionally aligned motor modules are preserved across slip directions. However, it is interesting that the existence of these shared synergies is driven by the choice of slip distance and slip speed. Additionally, we also found that the number of shared synergies varies with slip distance and speed. All the synergies reflected the effect of varying the slip distance, suggesting that the CNS primarily adjusts the required number of functionally aligned motor units depending on how much the object slips, rather than how fast it slips. However, the role of slip speed in sending the demand to the CNS to recruit or decruit the motor modules can not be completely ignored, as the number of shared synergies from motor modules of synergy 2 has shown a significant effect of variations in slip speed. Although similar studies on object slippage per se are scarce, we found several gait and balance recovery studies indicating that the CNS adjusts muscle synergies in response to slip perturbations [32]–[34]. Interestingly, no specific synergies were found associated with different slip directions, suggesting that CNS does not recruit distinct motor modules for upward and downward slip conditions. Our findings also suggest that several synergies were recruited differently based on the slip direction. In other words, the synergy from the upward slip direction was shared with multiple synergies in the downward slip direction, and this recruitment was found to be influenced by variations in both slip distance and slip speed. Such shared synergies can be interpreted as the same functional group of muscles being reconfigured into multiple direction-specific modules. These instances may indicate the existence of combined synergies, suggesting a flexible reconfiguration of shared synergies across different slip directions. However, it also raises an important question: Do these combined synergies reflect adaptive neural flexibility that allows shared muscle groups to be reorganized into direction-specific modules in response to task demands?

### C. Limitations and Future Work

Although the study offers unique insights into how the CNS modulates synergies during object slippage, certain limitations associated with the experimental protocol need to be acknowledged. The duration of the experiment was approximately 3 hours, which would have fatigued the muscles, and the muscle synergies have been shown to be affected by the muscle fatigue [35], [36]. Although these studies report that muscle fatigue affects the motor primitives, not the motor modules, the analysis of shared and specific synergies may not have a significant impact. However, extending this analysis to determine combined and pure synergies would also require analyzing motor primitives. Future work should either incorporate the dynamic synergy models [37] or the integrated analysis of motor modules and motor primitives to investigate how neural modules evolve during slips and how these synergistic patterns vary with variations in stimuli. Understanding these mechanisms would strengthen the development of rehabilitation techniques and the control strategies for prosthetic devices.

## V. Conclusion

This research work provides a unique insight into the dependence of muscle synergies on the slip characteristics. The findings from the study reveal that the existence of shared muscle synergies in the upward and downward slip conditions is governed by the choice of slip distance and slip speed. Additionally, the recruitment or decruitment of the functionally aligned motor modules by the CNS is also governed by the choice of slip parameters. These findings pave the way for developing a comprehensive understanding of the synergistic patterns that can ultimately be leveraged to build biologically inspired control architectures for bionic and rehabilitation robotic systems and advanced rehabilitation strategies.

