## Supplementary Information for "Task-Dependent Modulation of Upper-Extremity Muscle Synergies during Object Slippage"

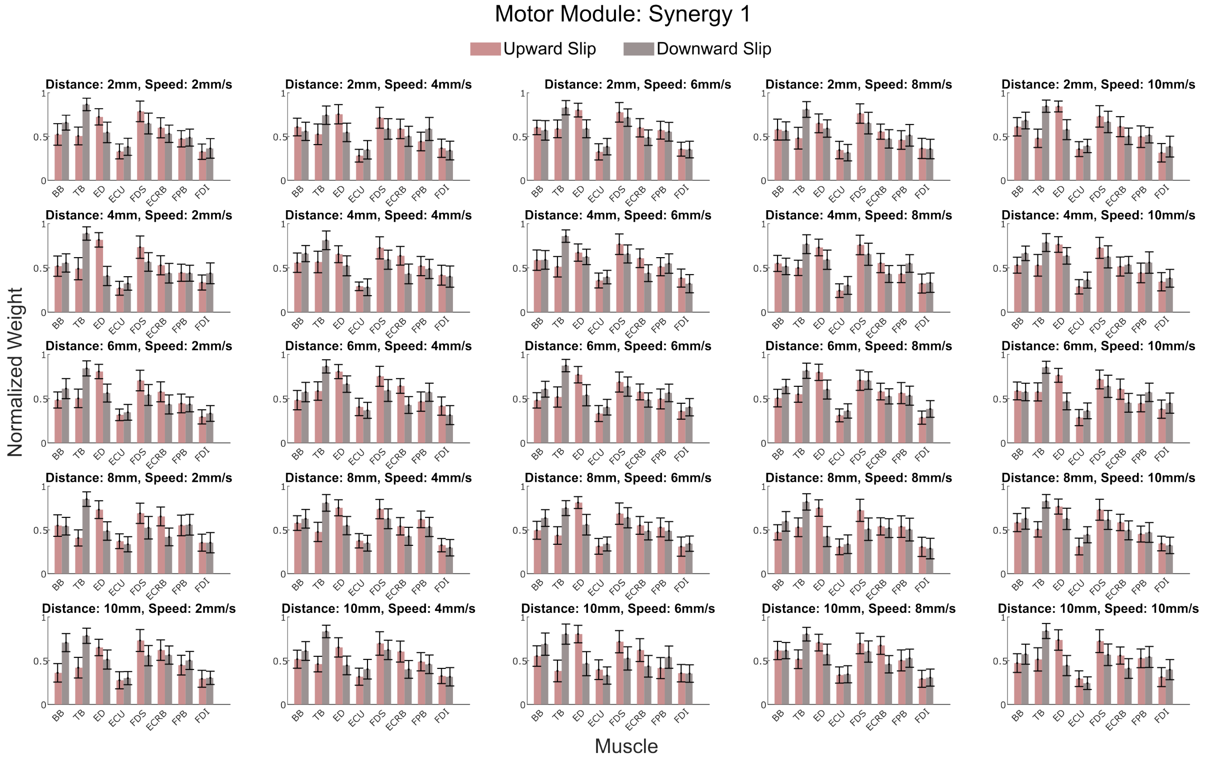


**Figure S1:** The average bar graphs for various combinations of slip distance and slip speed in the upward and downward directions corresponding to the motor modules of Synergy 1.

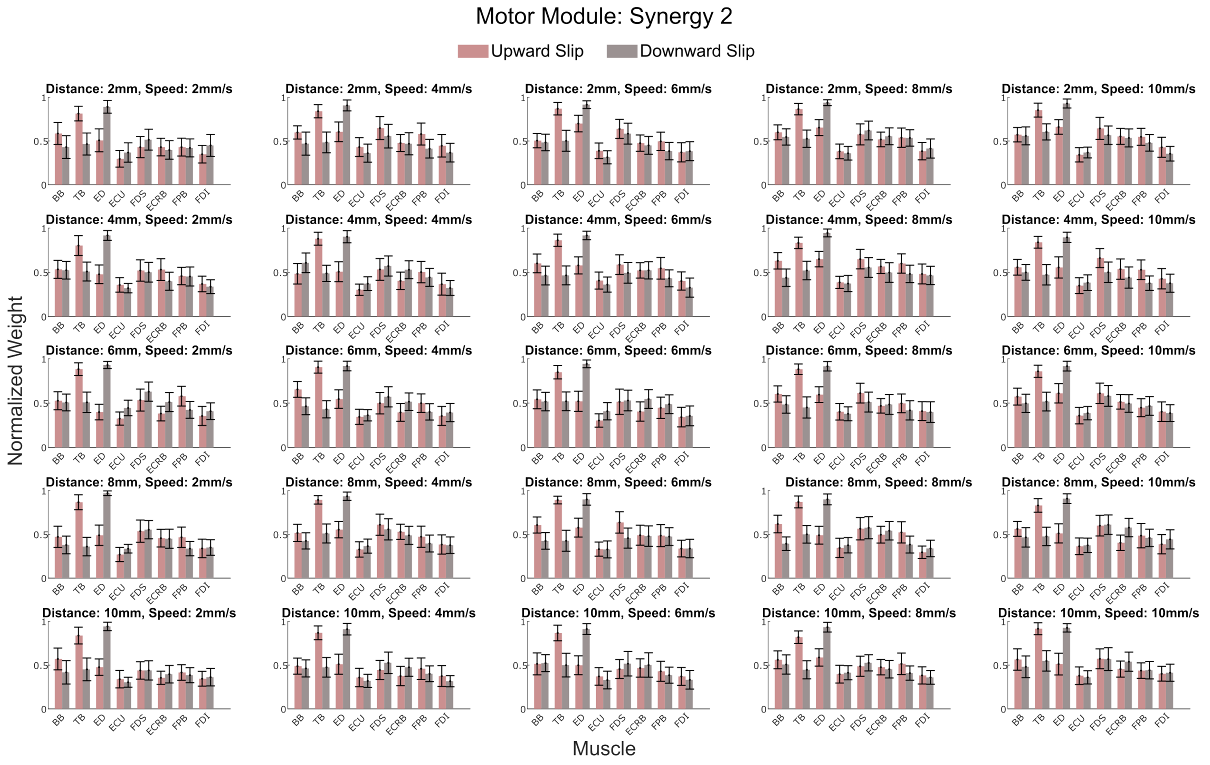


**Figure S2:** The average bar graphs for various combinations of slip distance and slip speed in the upward and downward directions corresponding to the motor modules of Synergy 2.

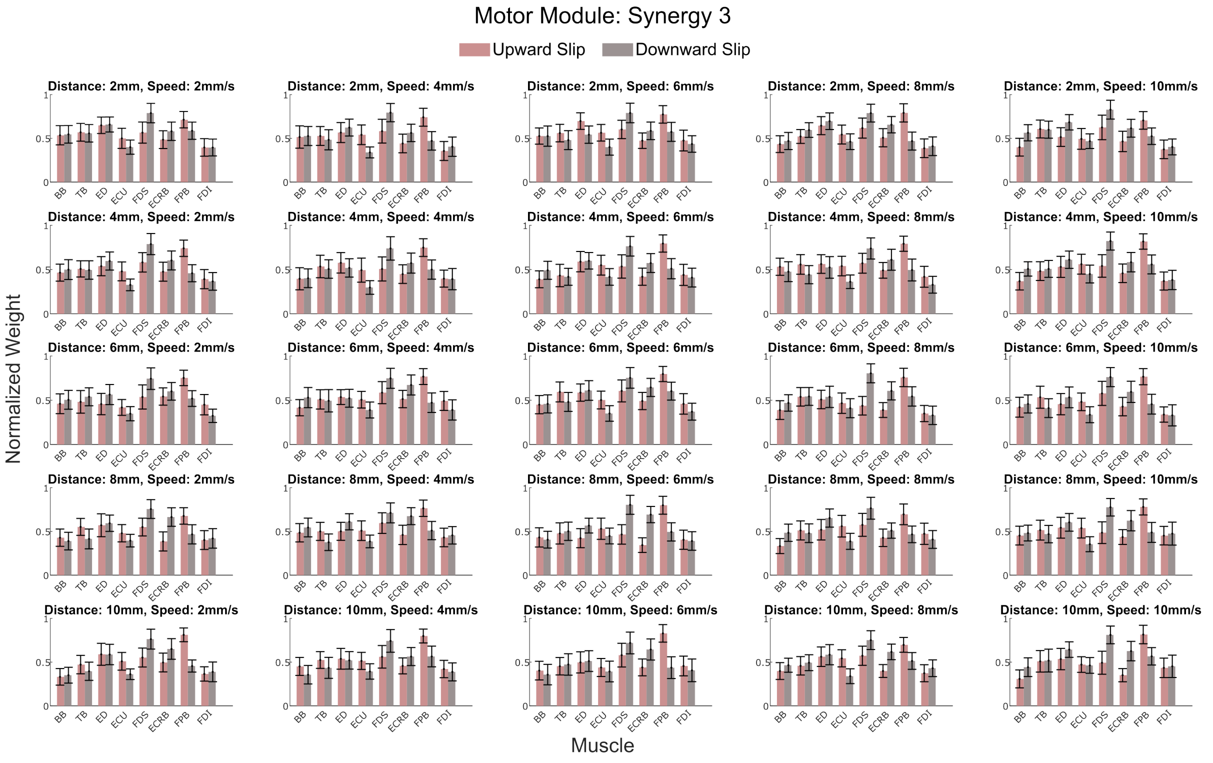


**Figure S3:** The average bar graphs for various combinations of slip distance and slip speed in the upward and downward directions corresponding to the motor modules of Synergy 3.

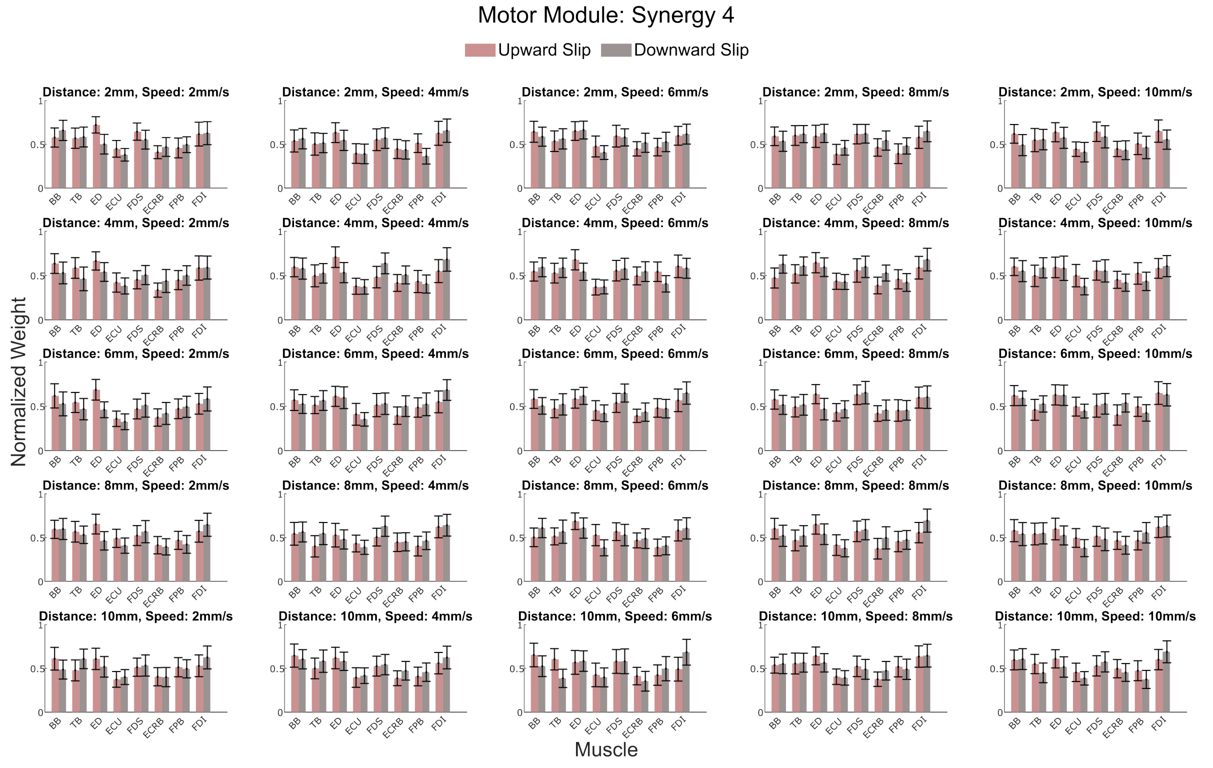


**Figure S4:** The average bar graphs for various combinations of slip distance and slip speed in the upward and downward directions corresponding to the motor modules of Synergy 4.

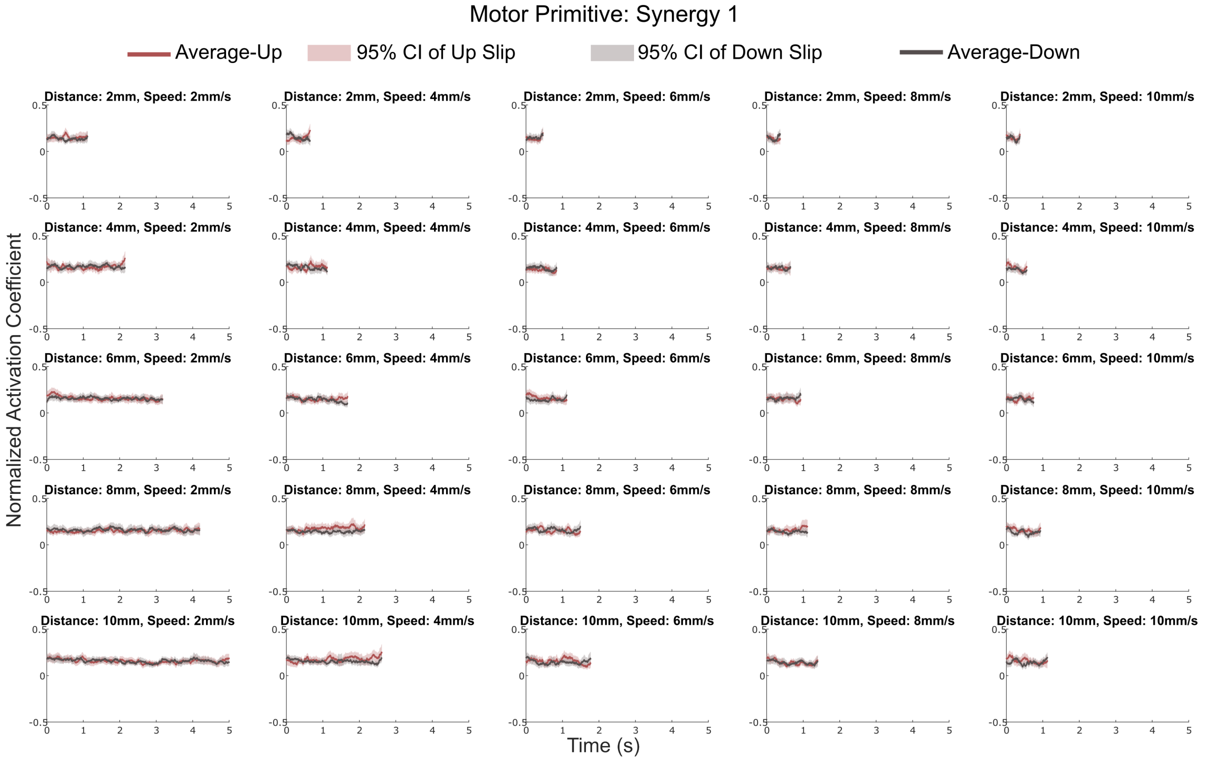


**Figure S5:** The average time series plots for various combinations of slip distance and slip speed in the upward and downward directions corresponding to the motor primitives of Synergy 1.

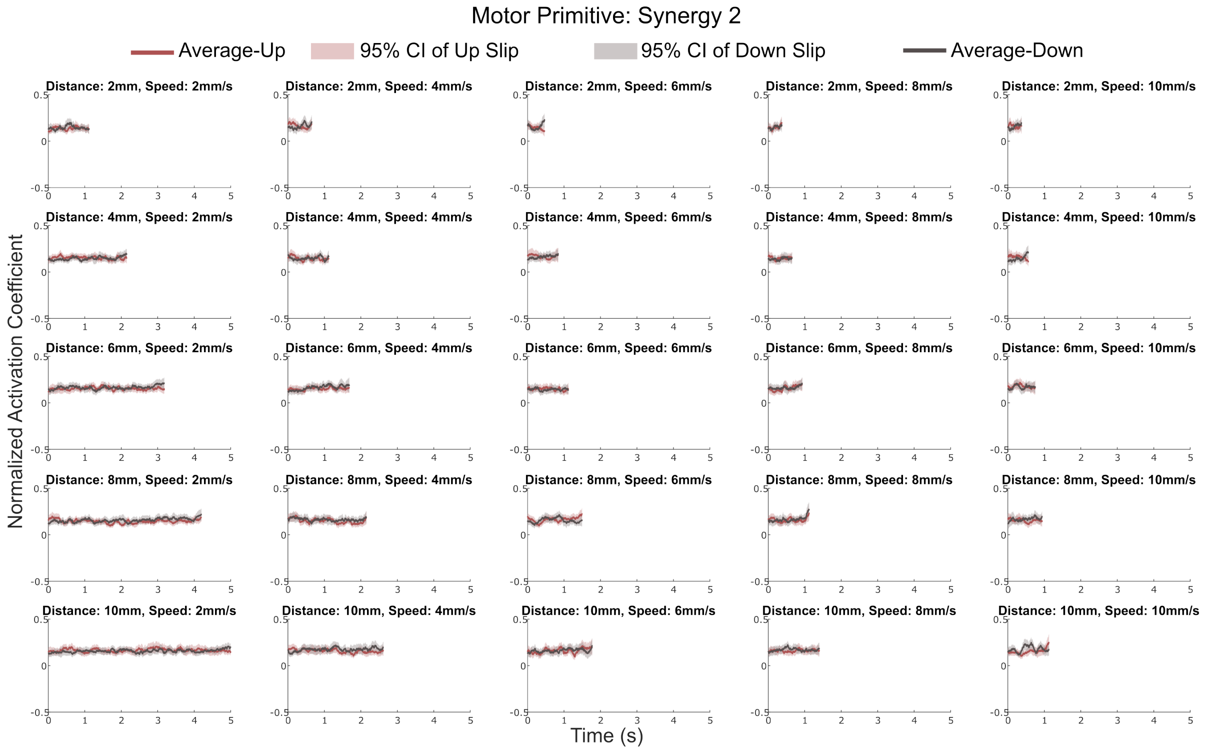


**Figure S6:** The average time series plots for various combinations of slip distance and slip speed in the upward and downward directions corresponding to the motor primitives of Synergy 2.

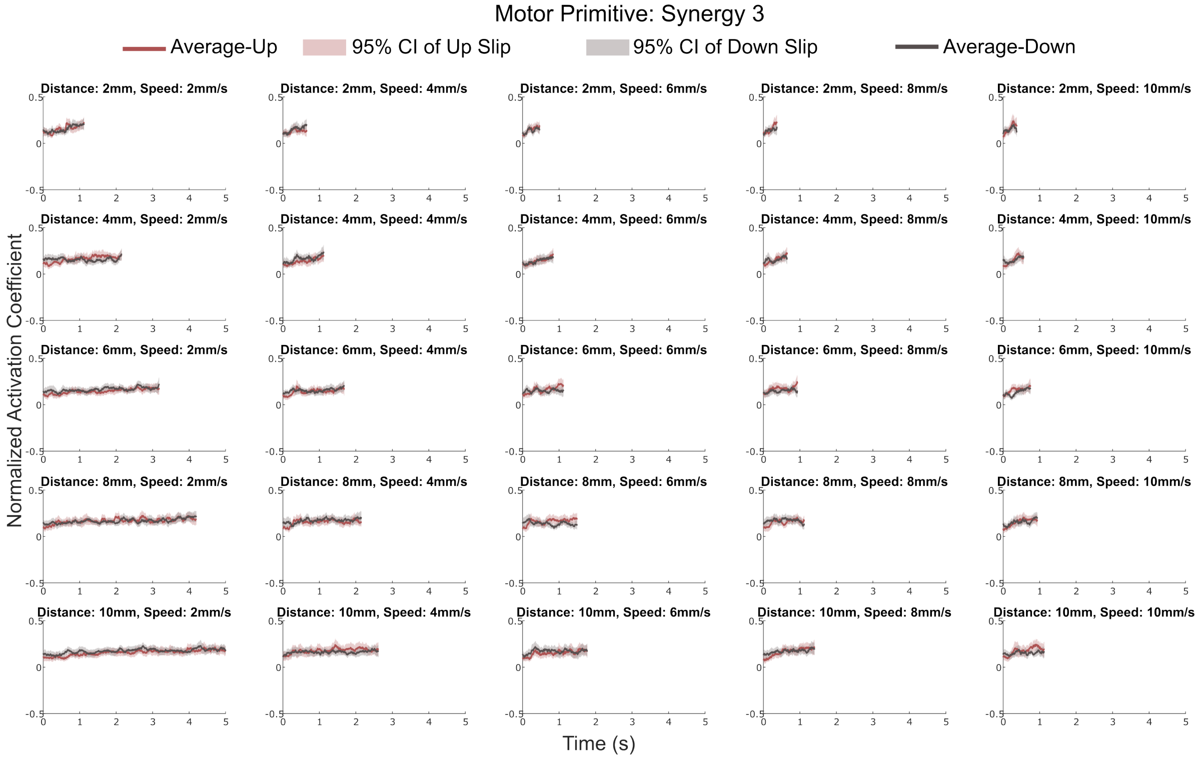


**Figure S7:** The average time series plots for various combinations of slip distance and slip speed in the upward and downward directions corresponding to the motor primitives of Synergy 3.

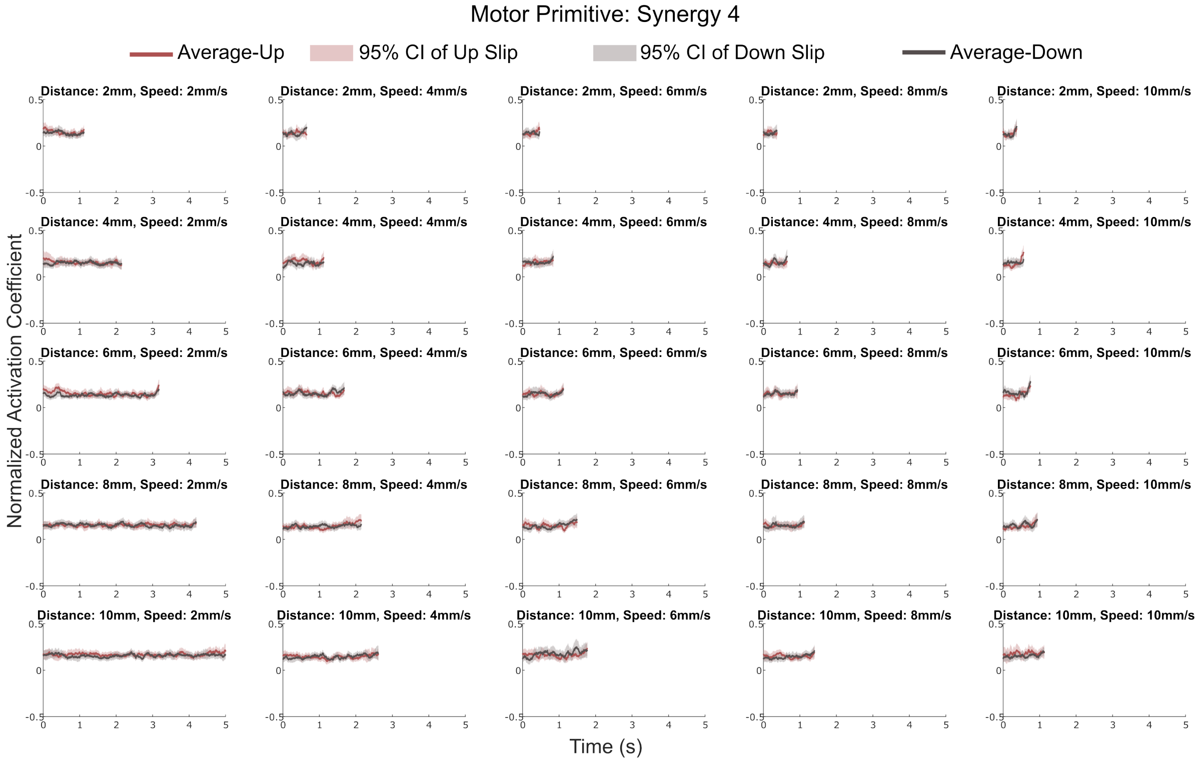


**Figure S8:** The average time series plots for various combinations of slip distance and slip speed in the upward and downward directions corresponding to the motor primitives of Synergy 4.
